# Time-resolved proteomics vs. ribosome profiling reveals translation dynamics under stress

**DOI:** 10.1101/087486

**Authors:** Tzu-Yu Liu, Hector H. Huang, Diamond Wheeler, James A. Wells, Yun S. Song, Arun P. Wiita

## Abstract

Many small molecule chemotherapeutics induce stresses that globally inhibit mRNA translation, remodeling the cancer proteome and governing response to treatment. Here we measured protein synthesis in multiple myeloma cells treated with low-dose bortezomib by coupling pulsed-SILAC (pSILAC) with high-accuracy targeted quantitative proteomics. We found that direct measurement of protein synthesis by pSILAC correlated well with the indirect measurement of protein synthesis by ribosome profiling under conditions of robust translation. By developing a statistical model integrating longitudinal proteomic and mRNA-seq measurements, we found that proteomics could directly detect global alterations in translational rate as a function of therapy-induced stress after prolonged bortezomib exposure. Finally, the model we develop here, in combination with our experimental data including both protein synthesis and degradation, predicts changes in proteome remodeling under a variety of cellular perturbations. pSILAC therefore provides an important complement to ribosome profiling in directly measuring proteome dynamics under conditions of cellular stress.

## Introduction

Dynamic changes in the cancer proteome control tumor growth, proliferation, metastasis, and response to the therapy. Targeting aberrant mRNA translation in cancer has recently garnered significant interest as a therapeutic strategy (Boussemart et al., 2014; Hsieh et al., 2012; Wolfe et al., 2014). Furthermore, a myriad of cellular stresses, including exposure to various chemotherapeutics, leads to global inhibition of protein synthesis and remodeling of the cancer proteome (de Haro et al., 1996; Walter and Ron, 2011).

A powerful new tool to measure gene-specific regulation of translation is ribosome profiling, the deep sequencing of mRNA fragments protected by actively translating ribosomes (Ingolia et al., 2009; Ingolia et al., 2011; Michel and Baranov, 2013). A central assumption of ribosome profiling is that indirect measurement of ribosome footprint occupancy on transcripts is directly reflective of true protein synthesis. While this assumption has been shown to be largely true in bacteria (Li et al., 2014a), the relationship between footprint occupancy and protein synthesis remains less clear in the more complex translational system of eukaryotes (Liu et al., 2016). Furthermore, using standard ribosome profiling approaches it can be difficult to capture global cellular changes in translational capacity (Ingolia, 2016), such as those which occur in response to drug therapy in cancer.

Here, we monitor the effects of low-dose bortezomib therapy on translation in multiple myeloma cells, a system of first-line therapy and hematologic cancer we have studied previously in the context of rapid apoptosis (Wiita et al., 2013). Proteasomal blockade by bortezomib is known to lead to endoplasmic reticulum stress due to accumulation of unfolded and misfolded proteins (Obeng et al., 2006; Vincenz et al., 2013). This stress triggers downstream signaling pathways that inhibit the translation of the large majority of mRNAs (Walter and Ron, 2011). Similar signaling to inhibit translation occurs in response to heat shock, DNA damage, and oxidative stress (Duncan and Hershey, 1984; Powley et al., 2009; Shenton et al., 2006).

Importantly, here we used pulsed-stable isotope labeling (pSILAC) approaches in combination with high-accuracy targeted quantitative proteomics to directly monitor the synthesis of new proteins in this system (Jovanovic et al., 2015; Schwanhausser et al., 2011). This design allowed us to compare ribosome footprint occupancy of transcripts to the directly measured synthesis of new protein molecules in a cancer therapy model. We found that during robust translation, before the onset of bortezomib-mediated translational repression, these two orthogonal measurements were well-correlated, providing important support for the assumption that ribosome footprint density is quantitatively reflective of protein synthesis in eukaryotes. We further developed a quantitative statistical model to describe protein synthetic rates, as derived from our proteomic data, against a background of general translational inhibition induced by prolonged bortezomib treatment. Under conditions of translational inhibition, we found that pSILAC methods were able to directly detect global alterations of translation not identified by standard ribosome profiling approaches. We further demonstrated that our model, incorporating experimental protein synthetic and degradation rates, could predict protein-level dynamics in response to different levels of stress-induced translational inhibition. These findings underscore the utility of pSILAC proteomics as a complementary method in studies of translational regulation to ribosome profiling, particularly under conditions of cellular stress.

## Results

We treated MM1.S multiple myeloma cells with the proteasome inhibitor bortezomib at a dose of 0.5 nM, well below the EC_50_ (~8 nM) for inhibition of proteasomal catalytic activity (Chauhan et al., 2005). We compared untreated cells (0h) to cells harvested at 6h, 12h, 24h, 36h, and 48h after treatment (Fig. 1A). At each time point we performed mRNA-seq and ribosome profiling as previously described (Wiita et al., 2013). We also harvested these unlabeled cells for quantitative proteomics using selected reaction monitoring (SRM). This method, employing a triple-quadrupole instrument, is used for high-accuracy quantitative measurements on cross-sample comparisons (Picotti and Aebersold, 2012).

**Figure 1.**
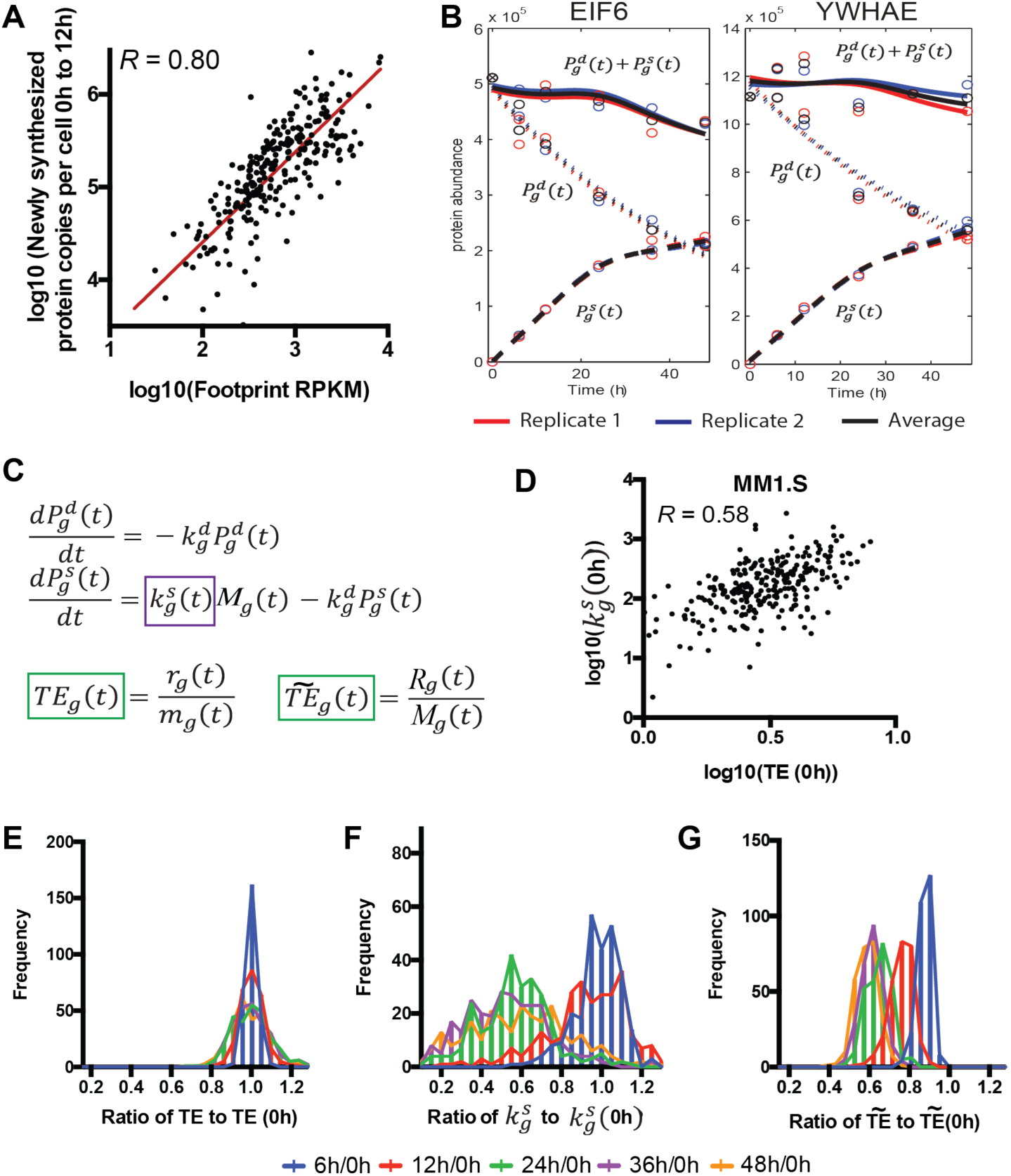
**Monitoring protein synthesis under low-dose bortezomib treatment. A.** Experimental design in MM1.S myeloma cells. **B.** Under 0.5 nM bortezomib there is little change in cell viability nor significant evidence of caspase activation. Values measured in duplicate +/− S.D. Normalized to 0h = 1 for cell viability (measured with Promega CellTiter-Glo assay), 0h = 0.1 for caspase 3/7 activity (measured by Promega Caspase 3/7-Glo assay). **C.** Selected Reaction Monitoring (SRM) proteomic assays on total protein demonstrate measurable increases in many proteins under low-dose (0.5 nM) bortezomib treatment that cannot be detected under rapid translational shutdown after high-dose (20 nM) bortezomib. Values +/− S.D. measured in technical duplicate.

Under these conditions we found that MM1.S underwent only minor changes in viability and caspase activation (Fig. 1B), unlike in our prior study at 20 nM bortezomib (Wiita et al., 2013). Total protein as well as total RNA and mRNA concentration per cell did not show any significant decrease over the time course (Fig. S1A), unlike under high-dose bortezomib (Wiita et al., 2013). We compared total protein abundance by SRM for a set of proteins that demonstrated increased transcript abundance after bortezomib exposure in our prior study (Wiita et al., 2013). We found that while these proteins were not detectably increased in the high-dose bortezomib setting, consistent with our prior results (Wiita et al., 2013), many of these proteins were indeed increased in the low-dose setting (Fig. 1C).

### Quantification of newly synthesized proteins

The above results cannot directly distinguish the contribution of new protein synthesis vs. existing protein degradation to the total protein abundance. Therefore, we next moved to a pSILAC approach. MM1.S cells were grown in “light” SILAC media supplemented with unlabeled L-lysine and L-arginine. At time 0h, cells were pelleted and resuspended twice in “heavy” SILAC media containing ^13^C_6_-^15^N_4_ arginine and ^13^C_6_-^15^N_2_ lysine and 0.5 nM bortezomib. Cells were grown for 48h under these conditions and harvested as above. Cell viability showed similar changes after treatment (Fig. S1B and 1).

We designed quantitative SRM assays measuring synthesis (“heavy” channel) and degradation (“light” channel) of 272 proteins in this cellular system. This analysis included monitoring at least two unique sequence peptides per protein, in technical duplicate, in both the light and heavy channels by SRM (Fig. 2A). SRM data were normalized across time points using the total intensity (light + heavy channel intensity) of a panel of “housekeeping” proteins that remain unchanged at the transcript level (see methods). While SRM has the advantage of consistent quantification of targeted peptides across all time points, a main drawback is the lower throughput compared to “shotgun” proteomic methods. Therefore, our analysis is necessarily limited to a subset of expressed proteins.

**Figure 2.**
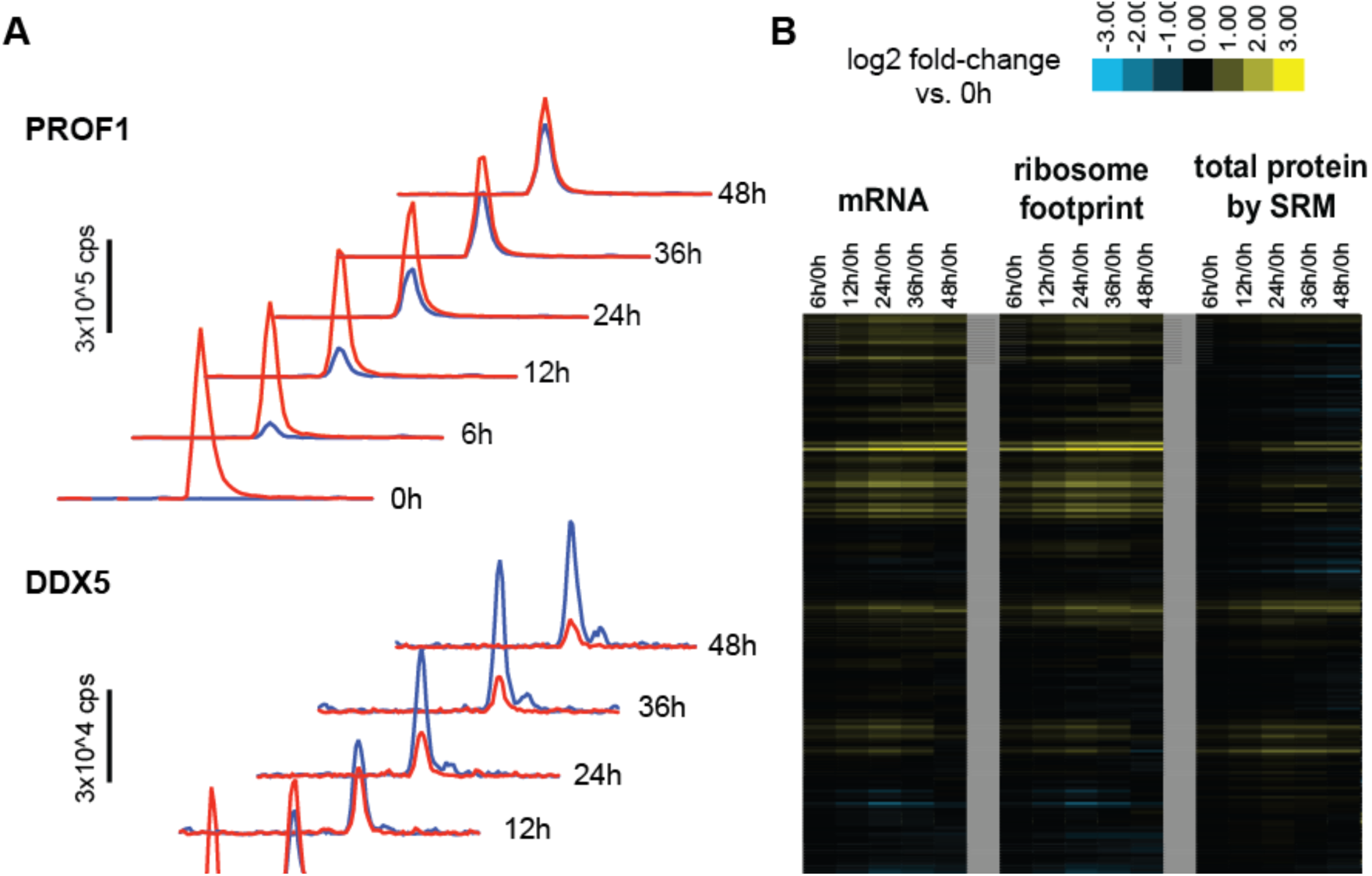
**Dynamics of transcript abundance and protein synthesis in response to low-dose bortezomib. A.** Example time-course SRM data for peptides from PROF1 and DDX5. Red traces = “light” channel intensity (degraded from baseline); blue traces = “heavy” channel intensity (newly synthesized post-SILAC pulse). Each trace represents added intensity of all monitored SRM transitions (four per peptide per channel). **B.** Relative mRNA abundance and ribosome footprint read density (ratio vs. 0h, in RPKM) move together over the timecourse whereas transcript-level changes are not seen as prominently as changes in protein abundance. SRM data here represents addition of all peptide intensities in light and heavy channels to measure total protein.

### Overall changes over the time course

We first compared the relative read density of transcripts identified by mRNA-seq and ribosome profiling across the time course to those found at baseline (untreated cells at 0h). We found that relative ribosome footprint density generally moves in concert with relative transcript abundance (Fig. S1H-I). The biological effects of low-dose proteasome inhibition were similar to those seen previously at high-dose (Wiita et al., 2013), with prominent upregulation of proteasomal subunits and downregulation of ribosomal subunits (Table S1).

In Fig. 2B, we compared the fold-change relative to 0hr of total abundance of the 272 proteins monitored by SRM across the time course to that of mRNA-seq and ribosome footprint read density on the corresponding transcript. While relative increases in mRNA drove increases in protein abundance, most protein-level increases were less prominent than transcript-level increases. Furthermore, downregulated transcripts did not lead to detectable decreases in protein abundance over 48h. This finding is consistent with those of others (Jovanovic et al., 2015; Schwanhausser et al., 2011) suggesting that high-abundance proteins, as we primarily monitored here, typically have long half-lives. These half-lives may be further extended by partial blockade of proteasomal degradation by bortezomib treatment (Fig. S3E).

### Correlation between protein copies and ribosome footprint densities

We next compared the amount of protein synthesis inferred from ribosome profiling to that measured by SRM. We first used the iBAQ approach (Schwanhausser et al., 2011) in biological duplicate on untreated MM1.S cells to estimate baseline protein copy number per cell (Fig. S2A-B). To ensure that these baseline copy numbers were of the correct order of magnitude, using quantitative Western blotting we verified protein copy number per cell for three representative proteins, spanning the range of estimated copy numbers per cell (~10^5^ to ~10^7^) for the majority of proteins included in the SRM assay (Fig. S2C-D). Using the heavy-channel SRM intensity, representing newly synthesized proteins, and extrapolating from baseline protein copies per cell, we estimated the number of protein copies per cell synthesized between the 0h and 12h time points, when cellular protein synthesis appears largely unaffected by drug treatment (Fig. S2E-G). We compared these data to the average ribosome footprint density (in RPKM) across the 0h, 6h, and 12h time points (Fig. 3A). Importantly, we found a good correlation between ribosome footprint density and protein synthesis (Pearson *R* on log-transformed data = 0.80). A linear best fit to this data on a log scale resulted in a slope of 0.97 (95% confidence interval: 0.86 to 1.06). This strong correlation and linear fit with slope near unity in this eukaryotic system suggests that indirect measurement of synthesis via ribosome footprint occupancy for any gene indeed appears to quantitatively reflect absolute protein synthesis. However, the observed correlation is not perfect, requiring further exploration of potential causes of divergence between these two orthogonal measurements.

**Figure 3.**
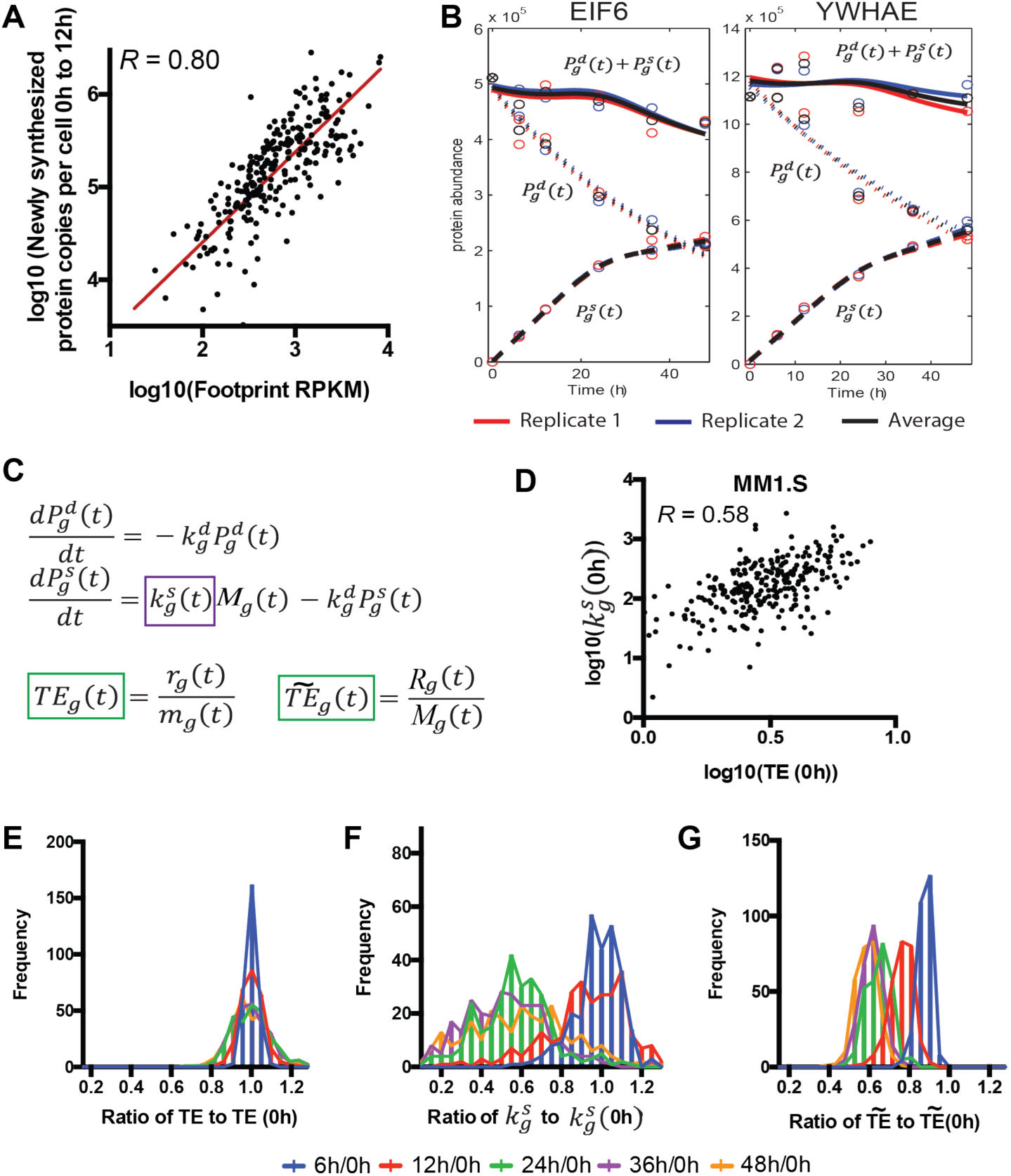
**Measuring and modeling protein synthesis and translational rate by pulse-chase proteomics and ribosome profiling. A.** Early in the time course, before any biochemical evidence of translational inhibition (Fig. S2), comparison of average ribosome footprint density on transcript coding sequence and newly synthesized proteins per cell, measured by SRM intensity and extrapolated based on iBAQ estimate of total protein copies per cell, show a strong correlation (Pearson *R*=0.80 on log-transformed data). Red line is line of best fit, with slope = 0.97 (95% confidence interval 0.86 to 1.06). Molecules synthesized per cell by SRM represent “heavy” protein copies at 12h minus “heavy” copies at 0h (see Methods); footprint data is average RPKM from the 6h, 12h, and 24h time points. **B.** Example plots from proteins EIF6 and YWHAE show proteomic data. Lines represent fits from model-fitting based on orthogonal splines with linearity constraints (see Methods). Solid lines = total protein; short dashed lines = degradation; long dashed lines = synthesis. **C.** Differential rate equation model describes protein abundance as a function of protein synthesis and degradation as measured from proteomic experiments and mRNA-seq. The primary free fitting parameter 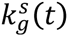 describes the number of proteins synthesized per transcript per unit time, 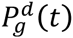 is abundance of “light” proteins, 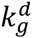 is protein degradation rate constant derived from single-exponential fit to light channel data, *M*_*g*_ is absolute transcript abundance as derived from mRNA-seq data, and 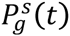 is abundance of newly synthesized “heavy” protein. The definition of translational efficiency from ribosome profiling literature is denoted by *TE* per gene: ribosome footprint read density *r*_*g*_(*t*) divided by mRNA-seq read density *m*_*g*_(*t*) (both in RPKM). The normalized translational efficiency is denoted by 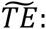 ribosome footprint read density normalized to mitochondrial footprints *R*_*g*_(*t*) divided by absolute transcript abundance *M*_*g*_(*t*). **D.** Baseline comparison of *TE* and 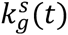 show that they are correlated, but imperfectly (Pearson *R*=0.58 on log-transformed data). *TE* (**E**), 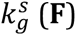 and 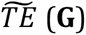 plotted across all 272 proteins across the time course, as a ratio to the value at 0h, shows no change in *TE* by standard analysis but similar changes in translational rate as measured by proteomics (**F**) and mitochondrial-corrected ribosome footprints (**G**).

### Mathematical modeling of protein synthesis and degradation

We therefore further explored the dynamics of protein degradation and production by using a system of differential equations. For each protein, we fitted the estimated number of “heavy” and “light” protein copies per cell, as well as the total protein abundance based on the addition of these two SRM intensities (Fig. 3B), using orthogonal natural cubic splines with linearity constraints to obtain functional forms, denoted by 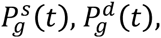 and 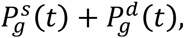 respectively (see methods). This enabled us to describe changes of protein abundance (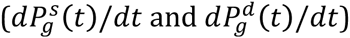 in terms of protein synthesis and degradation (Fig. 3C). For each corresponding transcript we also monitored the mRNA-seq read density (Mortazavi et al., 2008) and ribosome footprint density in RPKM.

To estimate the degradation rate constant 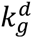 for each gene *g*, we found that a single-exponential fit well-described protein degradation for the included proteins. Other proteomic and deep sequencing data were fitted using the same approach described above (see Methods). The primary gene-specific free parameter in this analysis is 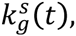 the translation rate parameter for gene *g* describing the number of protein molecules produced per transcript per unit time, thereby providing a proteomic-based measure of translational efficiency for each gene.Importantly, our model allows us to determine changes in 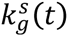 as a function of time, unlike in prior approaches describing a static 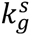 term (Schwanhausser et al., 2011).

### Comparison of inferred translational rate parameter from proteomics with translational efficiency from ribosome profiling

We can now directly compare 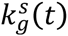 with a measure of translational efficiency (*TE*) used in the ribosome profiling literature, where *TE* is defined as the ratio of the relative ribosome footprint read density to the relative mRNA-seq read density (Ingolia et al., 2009; Ingolia et al., 2011). Using standard ribosome profiling analysis methods (see Methods), we observed little change in *TE* (Fig. 3E). This finding is in surprising contrast to changes found in 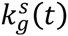 as measured by proteomics, where comparisons to 0h indicate a reduction in proteins synthesized per transcript across the time course (Fig. 3F).

We reasoned that our proteomic methods may be directly detecting global decreases in translational capacity induced by bortezomib treatment not captured by standard ribosome profiling approaches. To support this notion, polysome analysis by sucrose gradient centrifugation (Fig. S2E), incorporation of puromycin into nascent proteins (Fig. S2G), and dephosphorylation of the translation initiation factor eIF4E-binding protein 1 (4EBP1) (Fig. S2F) all supported a diminishment of translational capacity at time points after 12h, despite little decrease in global mRNA levels compared to baseline (Fig. S1A).

The inability of standard ribosome profiling approaches to detect global changes in translational capacity has been recognized previously (Ingolia, 2016). To address this issue, we used a recently described method of normalization incorporating ribosome footprints mapping to mitochondrially-encoded genes (ChrM), which are proposed to remain constant despite inhibition of cytosolic translation (Iwasaki et al., 2016). The normalized translational efficiency, denoted by 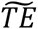 (see Methods), demonstrates qualitative agreement between changes in global translational capacity as measured by both ribosome profiling and proteomics across the time course (Fig. 3F-G). At 48h we measure a median ~40% decrease in translational efficiency of measured transcripts by both methods, albeit with greater variance in the proteomic measurement. This decrease in translational efficiency also appears consistent with biochemical measurements (Fig. S2E-G).This finding highlights that monitoring protein synthesis by mass spectrometry can directly confirm global changes in translational capacity.

### Modeling changes in global protein synthetic capacity

We note that the normalization method above may be limited by the low number of ribosome footprint reads mapping to chrM (Fig. S1J) or the long duration of low-dose bortezomib treatment. We therefore developed an algorithm to computationally estimate the changes in global protein synthetic capacity. A scaling function *G*_r_(*t*) (common to all genes) was incorporated into the system of differential equations. This function *G*_r_(*t*) was used to correct *TE* to reflect changes in the global synthetic capacity in the cell (blue curve in Fig. 4B). We inferred *G*_r_(*t*) that optimizes a squared loss function based on proteomic data (see Methods). When our inferred *G*_r_(*t*) was included in simulations of low-dose bortezomib treatment, the resulting simulated protein synthesis dynamics were similar to those noted by puromycin incorporation (Fig. S2G-H).

**Figure 4.**
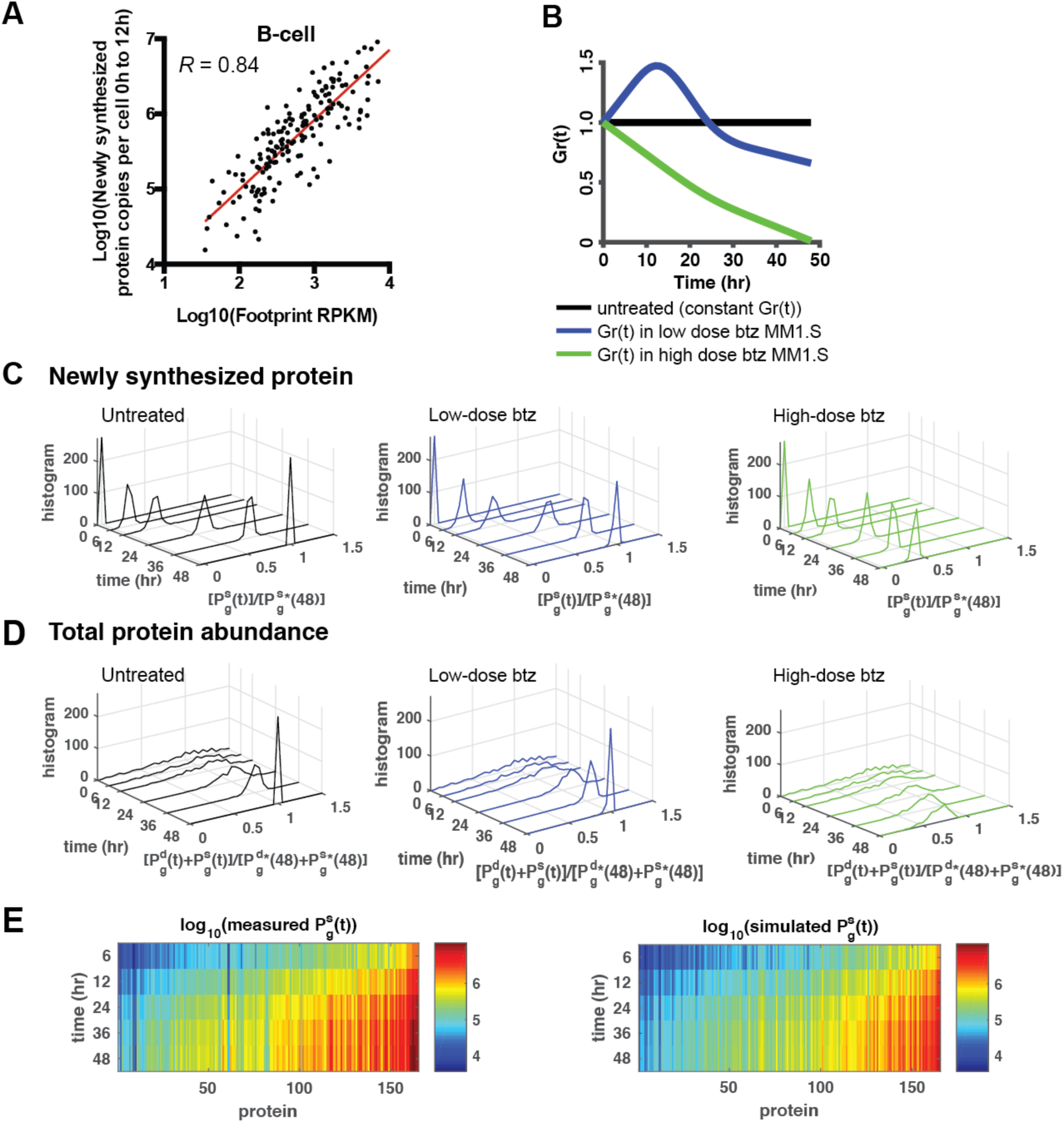
**A dynamic model predicts absolute protein synthesis under conditions of cellular stress. A.** Untreated Epstein Barr Virus (EBV)-immortalized B-cells also show a strong correlation (Pearson R on log-transformed data = 0.84) between absolute protein synthesis and footprint RPKM as in Fig 3A. Red line is line of best fit, with slope = 0.93 (95% confidence interval 0.84 to 1.02) **B.** Simulation of absolute protein copies synthesized as a function of different levels of translational capacity (denoted by *G*_*r*_(*t*)) in MM1.S, with inputs of iBAQ protein copy number, ribosome footprint density, mRNA-seq, *β* and 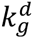 from MM1.S data. Three conditions are considered: *G*_*r*_(*t*) being constant, as in untreated cells; *G*_*r*_(*t*) varying as found in low-dose bortezomib (btz)-treated MM1.S; and *G*_*r*_(*t*) decreasing towards zero, as in high-dose bortezomib-treated MM1.S (Wiita et al., 2013) **C.** The simulated heavy channel protein abundance 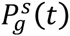 normalized by the heavy channel protein abundance at 48 hr under constant *G*_*r*_(*t*), denoted by 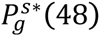 in MM1.S. Under high-dose btz, protein synthesis is strongly curtailed, consistent with Wiita et al. (2013). **D.** The simulated total protein abundance 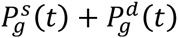 normalized by the total protein abundance at 48 hr under constant *G*_*r*_(*t*), denoted by 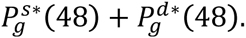 High-dose btz again shows a significant decrease in total protein abundance. **E.** SRM pSILAC-measured absolute protein copy number synthesis per B-cell (*left*) compared to simulated estimates of total absolute proteins synthesized in untreated B-cells over 48 h using quantitative model (*right*), with inputs of iBAQ protein copy number, ribosome footprint density, and mRNA-seq measured in B-cells and *β* and 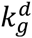 from MM1.S data, show good agreement.

### iBAQ replicate error partially explains noise in correlation between proteomic and ribosome profiling

We also investigated the correlation at baseline between translational efficiency and 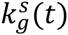 across proteins (Fig. 3D). We first estimated a multiplicative constant, *β*, that, when applied to all genes, largely reconciles the *TE* measured from ribosome profiling with 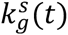 estimated from proteomic experiments (see Methods).A major question is whether the discrepancy of the fit between *TE* and 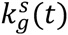 for different genes represents real biology (i.e. gene-specific translational regulation at the post-translational level, only detectable by proteomics) or systematic biases in one or both methods. Of note, our quantitative model relies on absolute protein copy number estimates from the iBAQ method (Schwanhausser et al., 2011). While the proteins included in our targeted SRM assay on MM1.S showed high reproducibility by iBAQ (Fig. S2B), simulations suggest that even this limited iBAQ replicate error could account for over a third of the residual variance of the correlation presented in Fig. 3A (Fig. S4B). To evaluate further potential sources of error, we examined whether accounting for annotated transcript isoforms could improve the correlation between footprint and proteomic data, but found only minor improvements (Fig. S4D).

### Predicting changes in protein synthesis

To further investigate applications of our model, we obtained a similar dataset of pSILAC proteomics paired with baseline mRNA-seq and ribosome profiling in untreated Epstein Barr Virus (EBV)-immortalized B-cells (Fig. 4A and Fig. S3). Importantly, the relationship between absolute protein synthesis and footprint RPKM (Fig. 4A) is also strong in this setting (Pearson *R* on log-transformed data = 0.84). A linear fit to this data on a log scale results in a slope of 0.93 (95% confidence interval 0.84 to 1.02). Again this strong correlation and linear relationship indicates that ribosome footprint occupancy is quantitatively reflective of absolute protein synthesis in a different cell type and without any drug perturbation.

Another direct application of our model is predicting absolute protein synthesis and abundance under different global levels of translational inhibition. Using our estimated values of 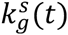 in combination with baseline mRNA-seq and ribosome profiling data, we predicted proteome remodeling under three different functions of *G*_r_(*t*) in MM1.S cells (Fig. 4B).Our model predicted significantly reduced absolute protein synthesis under conditions of strong translation inhibition (Fig. 4C and Fig. 4D), consistent with that found in our prior study of high-dose bortezomib (Wiita et al., 2013). We also predicted the “heavy” (newly synthesized) protein copy number in B-cells, using inputs of iBAQ protein copy number, ribosome footprint density, and mRNA-seq measured in untreated B-cells, and *β* and 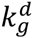 estimated from MM1.S data (Fig. 4E *right*), and found similarity to the experimental pSILAC measurements of heavy protein copy number (Fig. 4E-*left*). Our results from Fig. 3A and 4A further support the notion that ribosome profiling data, in combination with biochemical knowledge of global translational inhibition, may be sufficient to predict changes in the proteome using our quantitative model.

## Discussion

Here, we directly measured protein synthesis and ribosome footprint density in the setting of cancer therapy. Ribosome profiling has become a widespread technique to assess translational regulation and protein synthesis. One important question about this technique, however, is whether the resulting data truly reflect protein synthesis and translational rate. A recent study in *E. coli* demonstrated that when compared with previously published absolute copy numbers per cell, extrapolated synthesis rates based on ribosome footprint density correlated very well (*R* = 0.84)), supportive of the notion that ribosome footprint density, as measured by ribosome profiling, is directly reflective of absolute protein synthesis, even in the more complex translational system of eukaryotes (Jackson et al., 2010; Kozak, 1999).

Others have compared the capture and analysis of nascently-translated proteins by mass spectrometry to ribosome profiling data and found weaker correlations (*R* = 0.98) (Li et al., 2014a).In our work, we also find a strong positive correlation between ribosome footprint density and absolute protein synthesis as measured by targeted time-resolved pSILAC (Fig. 3A (*R* = 0.80) and 4A (*R* = 0.66) (Zur et al., 2016). However, this “Punch-P” approach has significant disadvantages as an orthogonal quantitative validation of ribosome profiling data as it relies on incorporation of a chain terminating puromycin analog for enrichment. Such truncated polypeptides will likely be rapidly degraded, skewing abundances in the captured cohort. Furthermore, enrichment-based methods suffer from biases in differential protein capture on streptavidin beads and artifacts from non-specific binding. These limitations make it difficult to quantitatively compare ribosome profiling to Punch-P. In contrast, the pSILAC approach we take here, combined with high-accuracy targeted quantification, allows us to directly measure protein synthesis in a complex system in an unbiased fashion.

With these data, we find that noise in baseline absolute protein abundance using the iBAQ methodology (Li et al., 2014b; Wilhelm et al., 2014) strongly affects the correlation between proteomic and ribosome profiling data. Other sources of error in our comparison that remain to be investigated may relate to ribosome footprint sample preparation methods (Weinberg et al., 2016) or splice isoform-specific translational control (Floor and Doudna, 2016). Due to these limitations we cannot exclude the possibility that for some genes there is a divergence between ribosome footprint occupancy and true protein synthesis, despite the overall strong correlation between these measurements across the monitored genes.

The quantitative model we develop also allows us to determine a measure of translational efficiency 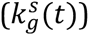 using proteomic data and compare this to translational efficiency as measured by ribosome profiling. Our results, with a Pearson *R* = 0.58 at baseline (Fig. 3D), are in line with that of a recent extensive time-course study of protein synthesis in murine dendritic cells (Jovanovic et al., 2015). They performed ribosome profiling at the baseline timepoint alone and also found a similar correlation (*R* = 0.5) between *TE* from ribosome profiling and 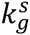 measured from shotgun proteomic data (Jovanovic et al., 2015). Given our findings (Figs. 3E-G), it appears that while proteomics may be able to broadly detect global changes in translational capacity, ribosome profiling may be more sensitive in determining translational efficiency changes for individual genes.

In the context of cancer therapy, our results here underscore that standard measurements of ribosome footprint density may not reflect absolute protein synthesis when global changes in translational capacity (i.e. the number of actively translating ribosomes) are present, whereas proteomics can more directly detect these changes (Fig. 3E and Fig. 3F). pSILAC combined with targeted mass spectrometry may therefore be an important method to orthogonally validate quantitative changes in translational rate found by normalization of ribosome profiling data under conditions of cellular stress (Andreev et al., 2015; Ingolia, 2016).

Furthermore, we develop a new quantitative model that can capture and predict dynamic changes in protein synthesis during cancer therapy. As we find a linear relationship between ribosome footprint density and absolute protein synthesis across genes, we suggest that with inputs of mRNA-seq, ribosome profiling, and absolute protein abundance estimates, in conjunction with biochemical data to describe the degree of translational inhibition, our model will provide a new window to predict the remodeling of the cancer proteome in response to therapeutic perturbation, even in the absence of a full pSILAC dataset. In addition, this quantitative framework can readily be applied to any human cellular system exposed to cellular stress impinging on the translational machinery.

## Acknowledgements

This work was supported by a Post-doctoral Fellowship (DRG 11-112) and Dale Frey Breakthrough Award (DFS 14-15) from the Damon Runyon Cancer Research Foundation, NCI K08CA184116-01A1, and an American Cancer Society Grant #IRG-97-150-13 (to A.P.W.); and NSF CAREER Grant DBI-0846015, a Packard Fellowship for Science and Engineering, and Math+X Research Grant from the Simons Foundation (to Y.S.S.). Mass spectrometry was performed at the UCSF NIH-NIGMS Mass Spectrometry Facility directed by Dr. Alma Burlingame supported by NIH grants P41RR001614 and S10RR026662. We thank Jessica Lund at the UCSF Center for Advanced Technology for assistance with HiSeq sequencing.

## Methods

### Experimental model and subject details

MM1.S cells were acquired from ATCC with cell line identity confirmed by karyotyping and DNA microarray. EBV-immortalized human B-cells derived from normal donor cord blood were a kind gift of Dr. Markus Müschen (UCSF Dept. of Laboratory Medicine) with normal diploid genome confirmed by karyotyping.

### Cell culture and drug treatment

MM1.S cells were grown in suspension to 1 × 10^6^ cells/ml in RPMI-1640 media with 10% FBS. EBV-immortalized B-cells were grown in suspension to 1 x 10^6^ cells/mL in RPMI-1640 media with 20% FBS. For initial MM1.S experiments (Fig. 1), bortezomib (LC Laboratories, Woburn, MA, USA) 20 µM stock solution in sterile-filtered phosphate buffered saline (PBS) was simultaneously added to a final concentration of 0.5 nM to flasks each containing 90 × 10^6^ cells (PBS only added to control sample). At the indicated time point cells were separated into aliquots for each experimental approach (15 × 10^6^ cells in duplicate for each of mRNAseq, proteomics, and ribosome profiling). Cells for ribosome profiling alone were incubated at 37°C for 1 min with 100 µg/mL cycloheximide (Sigma-Aldrich, St. Louis, MO, USA). All cells were pelleted by centrifugation, washed in PBS (PBS + 100 µg/ml cycloheximide for ribosome footprint samples), pelleted by centrifugation again, and flash frozen in liquid N_2_, then stored at −80°C. Cell viability and caspase activity were assessed by Cell-Titer Glo and Caspase-Glo (Promega, Madison, WI, USA) assays per manufacturer protocol, respectively. For pulsed-SILAC experiments, MM1.S cells were grown for 6 cell doublings in SILAC RPMI media depleted of arginine and lysine (Thermo Fisher Scientific, Waltham, MA, USA), dialyzed FBS (Life Technologies, Carlsbad, CA, USA) supplemented with unlabeled L-lysine (70 mg/L) and L-arginine (40 mg/L) (Sigma-Aldrich). EBV-immortalized B-cells were similarly grown for 6 cell doublings in SILAC RPMI media supplemented with “medium” (4,4,5,5-D_4_) lysine and ^13^C_6_ arginine. For both cell lines, at time 0h, 100 x 10^6^ cells at a density of 1 × 10^6^ cells/ml were pelleted by centrifugation and resuspended in SILAC RPMI media supplemented with “heavy” ^13^C_6_-^15^N_2_ lysine (70 mg/L) and ^13^C_6_-^15^N_4_ arginine (40 mg/L) (Cambridge Isotope Laboratories, Andover, MA, USA). Cells were harvested at the indicated time points, washed once in PBS, and stored at −80°C until sample analysis. For MM1.S cells, ribosome profiling, mRNA-seq, and proteomics were performed at each time point. For B-cell analysis, proteomics were performed at each time point, while ribosome profiling and mRNA-seq analysis were performed in biological duplicate on the baseline sample alone as the cells are not perturbed during the time course.

### Measurement of total RNA, mRNA, and protein

Total RNA was extracted either by Trizol (Life Technologies) per manufacturer protocol or using QIAgen RNeasy kit (QIAgen, Germantown, MD, USA). mRNA was further purified from isolated total RNA by poly(A) separation using Oligo (dT)_25_ Magnetic Beads kit (New England BioLabs, Ipswich, MA, USA) per manufacturer protocol. Total RNA and mRNA concentration was measured either by NanoDrop ND-1000 UV-Vis spectrophotometer (Thermo Fisher) or QuantiFluor RNA assay (Promega, Santa Clara, CA, USA). Total protein was isolated by lysis in either 8M Urea buffer for proteomics (see below) or RIPA buffer (EMD Millipore, Billerica, MA, USA) for immunoblotting and concentration measured using BCA assay (Thermo Fisher Scientific).

### Ribosome profiling and mRNA-seq

Ribosome profiling and mRNA-seq samples were prepared and analyzed as in our prior study (Wiita et al., 2013). Briefly, harvested cell pellets for ribosome profiling were suspended and lysed in 500 µl ice-cold polysome lysis buffer (20 mM Tris, pH 7.4, 250 mM NaCl, 15 mM MgCl_2_, 1 mM dithiothreitol, 0.5% Triton X-100, 24 U/ml Turbo DNase (Ambion, Austin, TX, USA), and 100 µg/ml cycloheximide). Lysate was clarified by centrifugation and RNase I 100 U/μl (Ambion) was added to digest polysomes to monosomes. Digested samples were then loaded onto a 1 M sucrose cushion and pelleted by centrifugation for 4 hr at 70,000 rpm. The pellet was resuspended in Trizol and RNA isolated per manufacturer protocol. RNA was separated by gel electrophoresis on a 15% TBE-Urea gel (Life Technologies) and gel fragments extracted corresponding to ∼25-35 nt in size. RNA was extracted from gel as in (Ingolia et al., 2011), by disrupting gel slices with centrifugation through a needle hole between 0.5 mL microfuge tube nested in a 1.5 mL microfuge tube. The gel was extracted in RNase-free water for 10 min at 70°C. The eluate was recovered by loading slurry onto a Spin-X column (Corning 8160) and centrifuging to recover eluate in collection tube. RNA was then precipitated from the filtered eluate by adding sodium acetate to a final concentration of 300 mM as a coprecipitant, followed by at least one volume of isopropanol. Precipitation occurred at −20°C overnight, RNA was pelleted and centrifuged for 45 min at 20,000 x g, 4° C. The supernatant was discarded and the RNA pellet was air dried, then resuspended in 10 μl 10 mM Tris (pH 7.0).

Harvested cell pellets for mRNAseq were isolated by Trizol and total RNA isolated per manufacturer protocol. Poly(A) mRNA was purified from the total RNA sample using poly-dT magnetic beads (as above) per manufacturer protocol. mRNA was fragmented in high pH buffer (50 mM NaCO_3_, pH 9.2) for 20 min at 95°C, then precipitated and separated by gel electrophoresis as above. mRNA fragments of 50-90 nt were extracted.

Both poly(A)-selected and ribosome footprint size-selected RNA samples were dephosphorylated, ligated to linker, and separated by gel electrophoresis as described previously (Ingolia et al., 2011). RNA was dephosphorylated with T4 DNA polynucleotide kinase (NEB M0201S), by resuspending RNA in 25ul 10mM Tris (pH 8.0), denaturing the fragments for 2 min at 75°C then equilibrating at 37°C and brought to a volume of 50 μl in 1X T4 polynucleotide kinase reaction buffer with 25 U T4 polynucleotide kinase (NEB M0201S) and 12.5 U Superasein (Thermo Fisher AM2694). This dephosyporylation reaction was incubated for 1 hr at 37°C and enzyme heat inactivated at 70°C for 10 min, then purified by precipitation as described above. Linker was ligated in a 20 μl reaction with dephosphorylated RNA, 12.5% w/v PEG 8000, 10% DMSO, 1X T4 RNA Ligase 2, truncated (NEB M0242L) reaction buffer, 20 U Superasein, 500 ng preadenylated miRNA cloning linker 1 (IDT), 200 U T4 RNA Ligase 2 (tr). This ligation was incubated at 37°C for 2.5 hr and the products were separated by gel electrophoresis and extraction as described above. Reverse transcription and cDNA library preparation were completed as in (Ingolia et al., 2011). Reverse transcription was carried out by preparing a reaction with RNA in 18 μl SuperScriptIII (Thermo Fisher 18080044) and 50 pmol oNTI-225 link1 primer (see key resource table). Reactions were denatured for 5 min at 65°C, equilibrated at 48°C with 2.0μl 1N NaOH and incubating 20 min at 98°C.Products were purified by gel electrophoresis and extracted as described above. Reverse transcription products were circularized with 20 μl CircLigase (Epicentre CL4111K) reaction. After circularization, subtraction of rRNA sequences was performed by subtractive hybridization using biotinylated oligos that reverse complement overabundant rRNA contaminants (see key resource table: oNTI309, 301r, 305r, 397hr, 298r, 303hr), by being suspended in 30 μl 2X SSC with 250 pmol total biotinylated subtraction oligos. The sample was denatured for 2 min at 70°C and transferred to 37°C. Hybridization was incubated for 30 min at 37°C.

Biotinylated oligos were removed by MyOne streptavidin C1 dynabeads (see key resource table), using 1mg magnetic beads. The rRNA-subtracted, circularized cDNA was used as a template for PCR amplication (see key resource table for amplification primers) using Phusion polymerase (NEB M0530S). Reaction products were then separated by gel electrophoresis as described above and DNA extracted using same procedure as above, with NaCl substituted as a co-precipitant. Extracted DNA was resuspended in 10 μl 10 mM Tris (pH 8.0) and expected library size verified using an Agilent Bioanalyzer 2100.

Sequencing was performed on an Illumina HiSeq 2500 using single end, 50-bp reads at the UCSF Center for Advanced Technology. Before alignment, linker sequences were computationally removed from the 3′ ends of raw sequencing reads.

STAR_2.4.0j was used to perform the alignments with up to one mismatch allowed. For footprint data only reads of length 25–36 nt (footprint length with cycloheximide (Ingolia et al., 2011)) were used for alignment. Reads were first aligned vs mitochondrially-translated genes; aligned reads were filtered. Next, all remaining reads were aligned vs human non-coding RNA and tRNA sequences; aligned reads were discarded. Finally remaining reads were aligned to human transcriptome reference (downloaded from http://www.gencodegenes.org/18.html Sep 2016) on reference genome GRCh37.

Software Ribomap (Wang et al., 2016) was used with default settings to assign multi-mapped reads to isoforms according to the mRNA abundance of each isoform. mRNA and footprint read density were calculated in units of reads per kilobase million (RPKM) to normalize for gene length and total reads per sequencing run.

Unsupervised hierarchical clustering was performed using complete linkage across mRNA, footprint, and translational efficiency data with uncentered correlation in Cluster 3.0 and visualized in TreeView. Where indicated, gene lists were analyzed by NIH DAVID resource (Huang et al., 2009) using default settings for included genes and interaction networks and human selected as species.

### Selected Reaction Monitoring proteomics

Frozen cell pellets were lysed by probe-tip sonication in buffer containing 8M Urea, 50 mM NaCl, and 100 mM Tris pH 8.0 supplemented with 1x HALT protease and phosphatase inhibitor cocktail (Thermo Fisher). Lysates were cleared by centrifugation at 16,500 x *g* for 10 min and protein concentration measured using the BCA assay. Lysate containing ~500 ug protein was diluted to 200 uL with lysis buffer. Disulfide bonds were reduced with 5 mM dithiothreitol and cysteines alkylated with 10 mM iodoacetimide. Lysate was diluted 1:6 with trypsin dilution buffer (100 mM Tris pH 8.0, 1 mM CaCl_2_, 75 mM NaCl). Sequencing grade modified trypsin (see key resource table) was added at an enzyme:substrate ratio of 1:25. Proteins were trypsin digested overnight with agitation at room temperature. Samples were adjusted to pH <3 with trifluoracetic acid and precipitate removed by centrifugation at 16,500 x *g* for 10 min. Tryptic peptides were desalted on SepPak C18 columns (Waters, Milford, MA, USA), evaporated to dryness on a vacuum concentrator, and stored at −80°C. For mass spectrometry analysis peptides were resuspended in 0.1% formic acid to a final concentration of ~0.2 μg/μL.

We previously developed targeted, label-free Selected Reaction Monitoring (SRM) assays for 152 proteins in our prior work in MM1.s myeloma cells treated with 20 nM bortezomib (Wiita et al., 2013). We applied this same assay to our samples as shown in Figure 1. Here we further developed new SRM assays to measure relative protein quantification in both the light and heavy SILAC channels (as in Fig. 2). For method development, data-dependent (or “shotgun”) proteomic data acquired on an LTQ Orbitrap Velos (Thermo Fisher) in HCD mode in our prior study was imported into the open-source software Skyline (v. 2.5) (MacLean et al., 2010) to build targeted assays consisting of parent ion and fragment ion “transitions” based on MS2 sequencing data. For heavy channel analysis, we used calculated increases in *m*/*z* of *y*-ion fragments to develop targeted methods. Two to four peptides per protein were targeted in initial method development. Proteins for analysis were primarily chosen based on high MS2 fragment intensity in shotgun data. Peptides were chosen having unique sequence identity for the targeted protein based on canonical sequence in Uniprot database.

All SRM analysis was carried out on an QTRAP 5500 (SCIEX, Framingham, MA) triple quadrupole mass spectrometer interfaced in-line with a nanoAcquity UPLC system (Waters) identical to that on the LTQ Orbitrap Velos (Thermo Fisher) on which the spectral library was acquired (Analytical column: BEH130 (0.075 × 200 mm column, 1.7 μm; Waters)). LC buffer A = 0.1% formic acid in water; buffer B = 0.1% formic acid in acetonitrile. We injected ∼1 μg of tryptic peptides from MM1.S cells onto the mass spectrometer with the following conditions: Direct sample loading at 3% B for 10 min after injection, a linear gradient from 3%-35% B over 80 min, an increase to 90% B over 5 min, then held for 5 min, then a decrease to 3% B for 10 min (total run time 110 min). Unit resolution was used at Q1 and Q3. A three second duty cycle time was used for all runs. For unscheduled runs a 10 ms acquisition time was used per transition. Multiple injections were used to test for all targeted peptides. Using data analysis in Skyline software, peptides were selected for further method development based on 1) the signal detection (above baseline) of at least 5 of 7 co-eluting transitions in both the light (0h sample used for method development) and heavy (48h sample used for method development) channels; 2) a retention time within 7 min of that acquired in the initial spectral library (acquired under the same chromatographic conditions); 3) fragment ion intensity of similar rank to that found in the initial spectral library.

Peptides chosen for further development were then limited to the four most intense transitions in both the light and heavy channels as found in unscheduled runs. A scheduled SRM method was developed with a retention time window of ±5 min. We then applied this scheduled method across multiple injections, with a minimum scan time per transition of 10 ms, at each time point, in technical duplicate. We ultimately chose to include in our pulsed SILAC analysis only those peptides which demonstrated detectable SRM intensity above background and at a consistent LC retention time at all time points in both the light and heavy channels (with the exception of heavy channel at 0h, where we expect to detect only background signal). Therefore, we ultimately included 733 peptides from 272 proteins in the analysis here for analysis of MM1.S cells. We applied this same method to EBV-immortalized B-cells and found that 165 of these proteins demonstrated sufficient signal-to-noise for analysis, likely based on differential protein expression between the two cell lines; these 165 proteins were used for all B-cell analysis.

Peptide intensity in each sample was measured as the sum of all transition peak areas for that peptide in each of the light and heavy channels (as measured by analysis in Skyline). Total peptide intensity was measured as the sum of the light and heavy channel intensities, and total protein intensity was measured as the sum of total intensities for all peptides from that protein. To normalize peptide concentration across samples, we used peptides derived from a set of high abundance proteins not expected to significantly change during the time course based on transcript-level data. We derived an index based on the geometric mean total intensity of peptides from these ‘housekeeping’ proteins (ENO1, KPYM, PPIA, FLNA, ACTB, TUBA1B) and scaled SRM intensity of all peptides in each channel based on the median value of this index. Corrected peptide intensity was averaged across injections for each sample.

### Western blots

For quantitative Western blots, 10 x 10^6^ untreated MM1.S cells were counted by taking the average of measurements from both manual hemocytmeter and automated cell counting using a Sceptre instrument (EMD Millipore). Cells were pelleted, washed 1x in cold PBS, and lysed in RIPA buffer (EMD Millipore) supplemented with 1x HALT protease and phosphatase inhibitors (Thermo Fisher) by probe-tip sonification. Protein concentration in lysate was measured by BCA assay. Recombinant proteins (GAPDH, Abcam; Vimentin, PeproTech; Bid, Sino Biological, see key resource table) at the manufacturer’s indicated concentration were used to generate a standard curve. Lysate and recombinant protein were separated on Mini-PROTEAN anyKd TGX gels (Bio-Rad, Hercules, CA, USA) and transferred to 0.45 μm PVDF membrane (EMD Millipore). Membranes were blocked using Odyssey blocking buffer (LI-COR, Lincoln, NE, USA) and probed with anti-GAPDH rabbit monoclonal antibody, anti-vimentin rabbit monoclonal, and anti-Bid rabbit polyclonal (Cell Signaling Technology, see key resource table) diluted at 1:1000 in Odyssey blocking buffer. Membranes were washed and blotted with infrared reporter-conjugated secondary antibodies and imaged on a LI-COR Odyssey system. LI-COR Image Studio software was used to quantify standard curve and lysate band intensity. Western blots for phospho- and total 4EBP-1 were performed as previously described with identical reagents (Wiita et al., 2013, see key resource table). All blots were completed in biological duplicate.

### Puromycin incorporation

The bortezomib treatment time course was performed in biological duplicate. One hour prior to each time point, 1 μM puromycin (Sigma-Aldrich) was added to 4.5 x 10^6^ cells at 1.0 x 10^6^ cells/mL. Cells from were allowed to incorporate puromycin for one hour, pelleted, washed in PBS, and lysed in RIPA buffer as above. 25 μg of lysate was separated by gel electrophoresis and transferred to PVDF membrane as above. Membranes were probed with a mouse anti-puromycin monoclonal antibody (KeraFast, see key resource table) and imaged on LI-COR Odyssey system.

### Sucrose density gradient

10x10^6^ MM1.S cells at each time point were lysed in 500 μL buffer containing 20 mM Tris pH 7.5, 50 mM NaCl, 5 mM MgCl_2_, 30% glycerol, 1% Triton-X, 20 U/mL SuperASEin (Ambion), 1 mM DTT, and 0.1 mg/mL cycloheximide. Lysate was loaded over a 10%-50% sucrose gradient, centrifuged at 35,000 x g for 3 hrs, and polysomes analyzed using a Gradient Station (BioComp, Fredericton, NB, Canada) with absorbance measured at 254 nm.

### Estimation of absolute protein copy number at baseline

We previously used the iBAQ method (first described in (Schwanhausser et al., 2011)) implemented in MaxQuant (Cox and Mann, 2008) to estimate protein copy number per cell at baseline in MM1.S cells. Here we identically analyzed unlabeled trypic peptides from a biological replicate of untreated MM1.S cells on an LTQ Orbitrap Velos mass spectrometer (Wiita et al., 2013). We estimated protein copy number per cell *ρ*_*g*_ based on the ion current assigned to each protein group *iBAQ*_*g*_ and scaled by a constant *σ*, such that *ρ*_*g*_ = *σ*[*iBAQ*_*g*_]. By incorporating the molecular mass *μ*_*g*_ of each protein, the total mass of protein per cell *μ*_*total*_, and Σ_*g*_ *ρ*_*g*_*μ*_*g*_ = *μ*_*total*_, we derived the scaling constant *σ*, and thereby estimated the protein copy number per cell *ρ*_*g*_. For incorporation into the quantitative model, we used the mean of the absolute protein copy number per cell from two iBAQ replicate analysis, denoted as 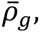 as the “0h” value for copies per cell.

This baseline quantity is related to the background corrected SRM intensity by 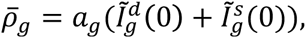 in which 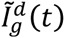 is the background corrected intensity of the light SRM channel, 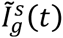 is the background corrected intensity of the heavy channel, and *a*_*g*_ is a gene specific constant. The background correction was conducted by subtracting the signal intensity in the heavy channel at 0h 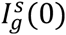 from all measurements, since no labeling had occurred, represented as 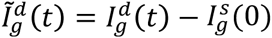 and 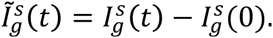 We extracted *a*_*g*_ using the background corrected SRM intensity and baseline absolute protein copy number per cell, and estimated protein copy number per cell at later time points in the heavy channel and light channel by 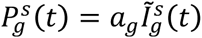 and 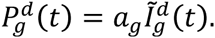

### Estimation of absolute mRNA copy number

We estimated mRNA copy number per cell *M*_*g,i*_(*t*) from mRNA-seq data using a method proposed previously (Schwanhausser et al., 2011; Wiita et al., 2013). Let *γ*_*g,i*_(*t*) represent the number of sequencing reads mapped to the transcript of gene *g*, isoform *i*, where *i* ∈ {1,…, *I*_*g*_} indexes the *I*_*g*_ isoforms of gene *g*. *l*_*g,i*_ represent the transcript length, and *T* (6.40×10^−16^ *mol*/*cell*) represent the total number of mRNA nucleotides per cell at baseline. These quantities can be related to the absolute mRNA copy number,

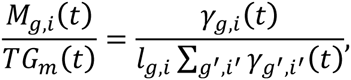

where *G*_*m*_(*t*) is the total mRNA nucleotide abundance at time t, normalized so that *G*_*m*_(0) = 1. Note that, up to a scale factor, the right hand side is the definition of *relative* mRNA abundance of gene *g*, isoform *i* (in RPKM), denoted by *m*_*g,i*_(*t*). Therefore, *M*_*g,i*_(*t*) = *c m*_*g,i*_(*t*)*G*_*m*_(*t*), where *c* is a constant. In what follows, we define 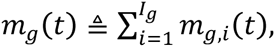 and 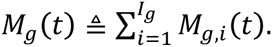

## Quantification And Statistical Analysis

Here we describe our quantitative model of protein synthesis dynamics and our inference procedure for parameter estimation.

### System of differential equations

We propose a system of differential equations to model the integrated longitudinal data of ribosome profiling, mRNA-seq, and pSILAC mass spectrometry. In what follows, we use the index *g* ∈ {1,2, …, *N*} to denote the gene or transcript ID, and use 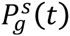 and 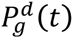 to denote, respectively, the newly synthesized protein abundance of gene *g* in the heavy channel and the degrading protein abundance of gene *g* in the light channel. The gene-specific degradation rate constant is denoted by 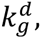, while the translational rate parameter for gene *g* is denoted by 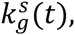 which is allowed to vary over time. Let *r*_*g*_(*t*)*G*_*r*_(*t*) denote the number of active ribosomes bound to each transcript of type *g* at time *t*, where 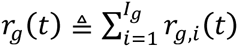 denotes the *relative* ribosome footprint abundance for transcript *g* (in RPKM), *r*_*g,i*_(*t*) denotes the *relative* ribosome footprint abundance for transcript *g* isoform *i* (in RPKM), *i* ∈ {1, …, *I*_*g*_} indexes the *I*_*g*_ isoforms of gene *g*, and *G*_*r*_(*t*) denotes the total number of active ribosomes. *G*_*r*_(*t*) is normalized so that *G*_*r*_(0) = 1. Ingolia *et al*. (Ingolia et al., 2009) defined “translational efficiency” as 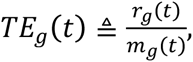 and we employ this definition.

In addition, we propose a new definition of translational efficiency that incorporates the ribosome footprints mapping to mitochondrially-translated genes. Let 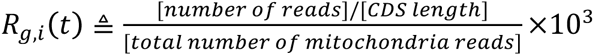 denote the normalized ribosome footprint abundance of gene *g* and isoform *i*, where *i* ∈ {1, …, *I*_*g*_} indexes the *I*_*g*_ isoforms of gene *g*. It is normalized by the total number of reads that mapped to mitochondrially-translated genes, representing reads per kilo base per mitochondria read. And let 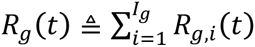 denote the summation across the isoforms of gene *g*. The new definition of translational efficiency becomes 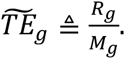 This definition incorporates the total cellular protein synthesis capacity *G*_*r*_(*t*) and total mRNA nucleotide abundance *G*_*m*_(*t*) into the calculation, which both can be time varying functions.

The proposed dynamical model of protein synthesis is a modification of the mass-action models for translation introduced earlier (Hargrove and Schmidt, 1989; Jovanovic et al., 2015; Schwanhausser et al., 2011; Wiita et al., 2013):

This model is a significant extension of our prior work (Wiita et al., 2013) in the following aspects: i) The pSILAC mass spectrometry technique enables us to extract the degradation rate constant 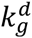 and thereby disentangle the degradation process from the synthesis process. ii) The relative ribosome profiling footprints measured in RPKM units are not sufficient to quantify how the absolute amount of footprints for each transcript varies over time. To address this problem, we incorporate a global function *G*_*r*_(*t*) (which we infer) that reflects the total cellular protein synthetic capacity (Eq. 3-2). An alternative solution is to use the mitochondrial footprints normalized translational efficiency 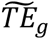 (Eq. 3-1). iii) Our modified model treats the translational rate parameter 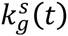 as a time varying function that depends on the density of ribosomes on the transcript. The major difference between the above model and that of Jovanovic *et al*. (Jovanovic et al., 2015) is that whereas their model utilizes only pulsed SILAC mass spectrometry data, we also incorporate the ribosome profiling footprint information into our model via Eq. 3-1 or Eq. 3-2.

Eq. 3-1

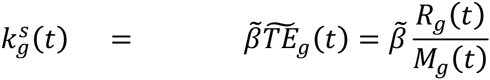

Eq. 3-2

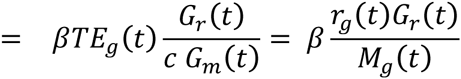

### Dilution effect resulted from cell growth

The number of cells in the low-dose bortezomib-treated MM1S data remains constant over time. Hence, 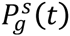 amd 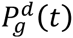represent respectively the heavy channel and light channel protein copies per cell. However, the untreated EBV-immortalized B-cells are growing, and the amount of light channel proteins per cell will be diluted due to cell division. We therefore scale the SRM intensities 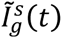 and 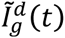 by the growth function *g*(*t*), and obtain the heavy channel and light channel protein abundance as 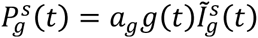 and 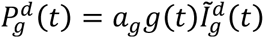respectively. This allows us to model the light channel protein abundance as an exponential function, discussed in the next section.

### Exponential and orthogonal natural cubic spline fitting

The solution to Eq. 1 is given by 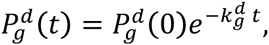 so fitting the observed degrading protein abundance with an exponential function yields an estimate of the degradation rate constant 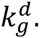. We employ the framework of functional data analysis in our study. In particular, measurements of 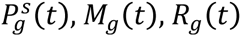 sampled at discrete time points are fitted with orthogonal natural cubic splines. Further details are provided below.

Eq. 1

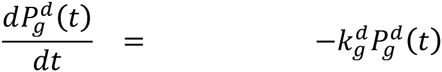

Spline is one of the widely used bases when approximating non-periodic functions (Ramsay and Silverman, 2005). We apply matrix factorization to the B-spline basis 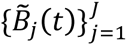 with degree 3 and three knots to construct an orthonormal basis 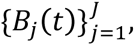 where *t* ∈ [*T*_1_, *T*_2_]. Let Σ be the Gram matrix of 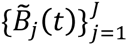 with the (*i, j*)-entry 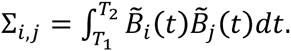 By matrix factorization, one can find an invertible transformation Λ such that Λ^T^Λ = Σ^−1^ and 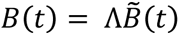 forms an orthonormal basis (Redd, 2012). Let 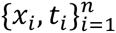 be a set of longitudinal data, where *x*_*i*_ is the measurement, *t*_*i*_ is the time when the measurement is observed, and *n* is the number of measurements. The data are approximated by a linear combination of the orthonormal basis functions 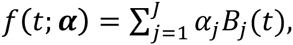 where ***α*** = (*α*_1_, …, α_J_). We adopt the penalized least-squares estimator to fit the function *f*(*t*; *α*) and utilize generalized cross-validation to decide the penalty parameter (Ruppert, 2002). Since the behavior of polynomials fit beyond the boundaries can vary erratically, we impose additional constraints such that *f*(*t*; *α*) is linear at *T*_1_ and *T*_2_, as in natural cubic spline (Hastie et al., 2009). The problem can be formulated as
Eq. 4

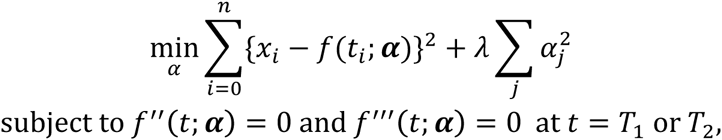

where *λ* is the penalty parameter, and *f″* and *f‴* denote the second and the third derivatives of *f*, respectively. This constrained optimization problem can be solved by the method of Lagrange multipliers. The Lagrangian of Eq. 4 can be written as

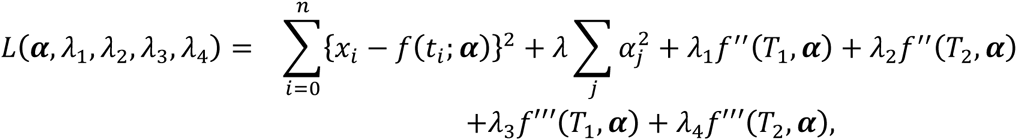

where *λ*_1_, *λ*_2_, *λ*_3_, *λ*_4_ are the Lagrange multipliers. The solution should satisfy ∇_*α*,*λ*_1_,*λ*_2_,*λ*_3_,*λ*_4__ *L*(*α*,*λ*_1_,*λ*_2_,*λ*_3_,*λ*_4_) = 0. For convenience, define *X* ≜ [*x*_1_ *x*_1_ … *x*_*n*_]^*T*^, 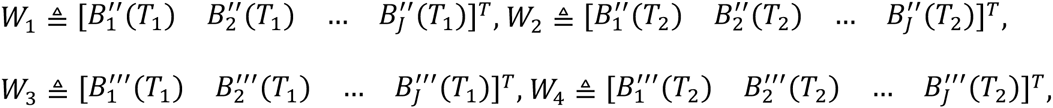
and

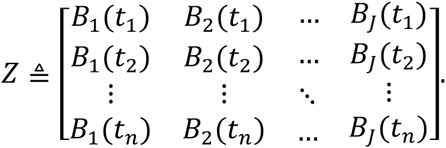

This then leads to a system of linear equations

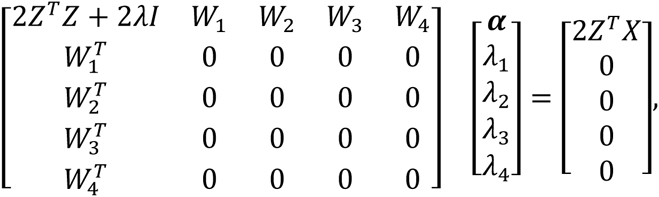

which can be solved by blockwise matrix inversion. Examples of the fitted functional form of 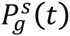 and 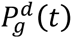 can be found in Figure 3B. Since the replicates of SRM measurements had high reproducibility, we use the average of them in the following analysis.

### Solving for parameters in the system of differential equations

We estimated the derivative of 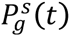 using its functional representation obtained from the mass spectrometry data, and fit 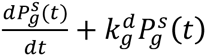 using degree-3 orthogonal natural cubic splines with 4 knots. Let 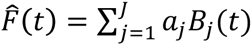 denote the resulting fit. Then, using Eq. 2 and Eq. 3-1, we solved for 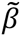 that minimizes the objective function

Eq. 2

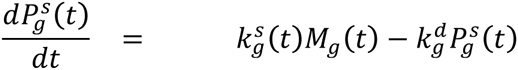

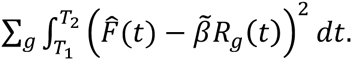

Fit *R*_*g*_(*t*) using degree-3 orthogonal natural cubic splines with 4 knots. Let 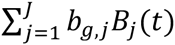 denote the fit. Then the objective function becomes

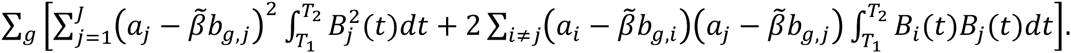

Since the basis is orthonormal, the objective function is simplified as a least-squares regression problem: 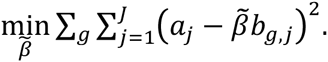

Similarly, using Eq. 2 and Eq. 3-2, we solved for *β* and the total cellular protein synthetic capacity *G*_*r*_(*t*) that minimizes the objective function

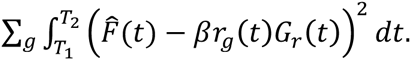

We achieved this by the following two steps:

1. Initialize *β* = 1.
2. Optimize with respect to *G*_*r*_(*t*):

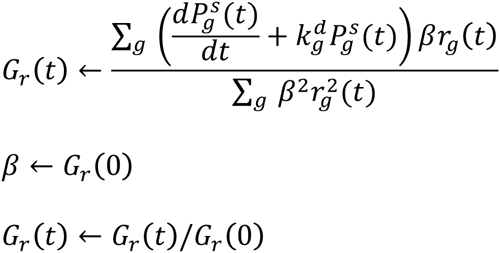

### Prediction of translational rate parameters and protein synthesis rates using ribosome profiling and RNA-seq measurements

To test whether *β* is a universal factor that applies both in the absence of and during exposure to bortezomib, we carried out leave-one-out prediction tests as follows. In testing gene *g*, we estimated *G*_*r*_(*t*) using the remaining *N* — 1 genes and estimated *β* using the protein measurement for *g* at time 0 (i.e., prior to the application of bortezomib). These parameters were then used together with ribosome profiling and RNA-seq measurements to predict the translational rate parameter 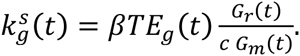 Then, the protein synthesis rate 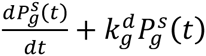 was predicted as 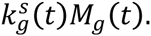 We assessed the performance of our predictions using the data for 6-48 hours, i.e., during exposure to bortezomib. The average performance in terms of the Pearson correlation coefficient and the relative mean absolute error are reported in Table S2, which illustrates that the error of our estimation and prediction procedures is comparable to the variation observed in experimental replicates.

Of note, Jovanovic et al. (Jovanovic et al., 2015) introduced the concept of the “recycling rate” in SILAC mass spectrometry experiments. This rate describes the incorporation of unlabeled amino acids into newly synthesized proteins after SILAC pulse. However, in our targeted SRM measurements, we can only measure fully “light” and fully “heavy” peptides. We cannot readily measure the infrequent mixed peptides, with both light and heavy residues in the same peptide, necessary for measuring the recycling constant γ. Therefore, we made a “shotgun” measurement at the 12h time point sample with an LTQ Orbitrap mass spectrometer using previous instrument and LC parameters (Wiita et al., 2013). We analyzed the results with Protein Prospector (prospector.ucsf.edu) using ^13^C6-^15^N4 arginine and ^13^C6-^15^N2 lysine as variable modifications. Of 372 peptides with at least one missed tryptic site (i.e. multiple lysines and/or arginines), and at least one labeled residue, only 8 showed evidence of mixed labeling. This result leads to a *γ*(12h) of 0.012, which has a negligible impact on measurement of protein synthesis using the model of Jovanovic et al. (Jovanovic et al., 2015). This result is also in line with their results, finding the recycling constant to have minimal effects after 12h of SILAC pulse (Jovanovic et al., 2015). Therefore, our measurements of protein synthesis, going up to 48h, are likely unaffected by not explicitly including the recycling constant in our model.

### Simulation of the biochemical experiment with puromycin incorporation

The biochemical experiment (Figure S2G) was conducted with 1 hour pulse of puromycin added at each time point prior to cell harvest. To test the accuracy of our proposed dynamical model, we performed a protein synthesis simulation using the parameters 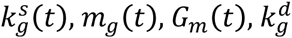 estimated from RNA-seq and mass spectrometry experiments. The initial condition was set such that 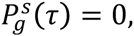 and the simulation was conducted to acquire the abundance after 1 hour, 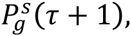 where *τ* ∈ {0, 6, 12, 24, 36, 48}. The simulation results qualitatively showed a similar pattern as the biochemical experiment (Figure S2H).

### Simulations of under- or over-estimation in iBAQ

We examined how much variation in 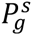 (Figure 3A and Figure 4A) can be explained by noise in iBAQ by performing simulations to model under- or overestimation in iBAQ values. Assume that the newly synthesized protein copies correlated perfectly with ribosome density, i.e. the red lines in Figure 3A and Figure 4A. We randomly generated scaling factors *s*_*g*_ according to the fitted normal distribution of the differences between the log-transformed iBAQ replicates (Figure S4A). Multiplying the ideal newly synthesized protein copies by these independent scaling factors, and repeating the process for 500 trials, we built a confidence interval for the abundance of newly synthesized protein copies (Figure S4B and Figure S4C). Then, we compared the average vertical offsets from the red line (Figure S4B and Figure S4C) in terms of MSE to the measured vertical offsets (Figure 3A and Figure 4A). Results showed that the iBAQ noise explained 36% and 23% of the deviation from the linear regression fits for MM1.S and B-cells respectively.

We further examined how the presence of transcript isoforms affected our estimation of these iBAQ noise statistics. We partitioned all 272 protein-transcript pairs monitored in MM1.S cells into two groups (Figure S4D), using the criterion: one dominant transcript isoform (>80% of RNA-seq read density on a single isoform, per paired-end RNA-seq analysis at www.keatslab.org/data-repository: HMCL66 Transcript Expression FPKM). The group that has one dominant transcript isoform has almost half (47%) of the deviation from the linear regression fits explained by iBAQ noise, whereas the other group has 35% of the deviation explained by iBAQ noise. This suggests that some other sources of error may exist, e.g., the difficulty of footprint alignment in the presence of transcript isoforms, which remains an open question.

